# Sepsis-induced NET formation requires MYD88 but is independent of GSDMD and PAD4

**DOI:** 10.1101/2024.07.29.605563

**Authors:** Hanna Englert, Chandini Rangaswamy, Mylène Divivier, Josephine Göbel, Irm Hermans-Borgmeyer, Uwe Borgmeyer, Kerri A. Mowen, Manu Beerens, Maike Frye, Reiner K. Mailer, Mathias Gelderblom, Evi X. Stavrou, Roger J. S. Preston, Stefan W. Schneider, Tobias A. Fuchs, Thomas Renné

## Abstract

Neutrophils are peripheral blood-circulating leukocytes that play a pivotal role in host defense against bacterial pathogens which upon activation, they release web-like chromatin structures called neutrophil extracellular traps (NETs).

Here, we analyzed and compared the importance of myeloid differentiation factor 88 (MYD88), peptidyl arginine deiminase 4 (PAD4), and gasdermin D (GSDMD) for NET formation *in vivo* following sepsis and neutrophilia challenge. Injection of lipopolysaccharide (LPS)/*E. coli* or the transgenic expression of granulocyte colony-stimulating factor (G-CSF), each induced NET-mediated lethal vascular occlusions in mice with combined genetic deficiency in *Dnase*1 and *Dnase1l3* (*D1*/*D1l3*^−/−^). In accordance with the signaling of toll-like receptors, *Myd88/D1/D1l3*^−/−^ animals were protected from the formation of lethal intravascular NETs during septic conditions. However, this protection was not observed during neutrophilia. It was unexpected to find that both *Gsdmd/D1/D1l3*^−/−^ and *Pad4/D1/D1l3*^−/−^ mice were fully capable of forming NETs upon LPS/E.coli challenge. Sepsis equally triggered a similar inflammatory response in these mice characterized by formation of DNA-rich thrombi, vessel occlusions, and mortality from pulmonary embolism, compared to *D1/D1l3*^−/−^ mice. Pharmacologic GSDMD inhibitors did not reduce PMA-stimulated NET formation in *ex vivo* models either. Similarly, neither Pad4 nor GSDMD deficiency affected intravascular occlusive NET formation upon neutrophilia challenge. The magnitude of NET production, multi-organ damage, and lethality were comparable to those observed in challenged control mice.

In conclusion, our data indicate that NET formation during experimental sepsis and neutrophilia is regulated by distinct stimulus-dependent pathways that may be independent of canonical PAD4 and GSDMD.

**Key points:**

- Sepsis triggers vaso-occlusive NET formation in *Dnase*1/*Dnase*1l3-deficient mice in a myeloid differentiation factor 88-dependent manner
- Peptidyl arginine deiminase 4 and gasdermin D are dispensable for NET formation in sepsis and neutrophilia models
- Myeloid differentiation factor 88, peptidyl arginine deiminase 4 and gasdermin D differ in their importance for NET formation *in vivo*

## INTRODUCTION

Neutrophils are white blood cells that play a central role in innate immunity. In bacterial infection, neutrophils are the first leukocytes to migrate to sites of inflammation in a process called chemotaxis. Upon activation, neutrophils secrete antimicrobial molecules, including myeloperoxidase (MPO) and neutrophil elastase (NE), which facilitate the eradication of infection. Additionally, activated neutrophils also release web-like structures termed neutrophil extracellular traps (NETs) (1). NETs are supramolecular aggregates of extracellular fibers, primarily composed of unfolded nuclear-derived double-stranded DNA (dsDNA) and histones. Together with granular mediators, they provide a physical barrier for pathogens and prevent the dissemination of infections (1). Although NETs are pivotal to host defense, unchecked excessive NET release (NETosis) or defective NET clearance can equally contribute to the development of inflammatory disorders including (auto)immune and thromboinflammatory diseases. Indeed, NETs have been demonstrated to prime immune cells to induce sterile inflammation or potentiate autoimmunity, for example by stimulating interferon responses (2, 3). The accumulation of platelets and coagulation factors within the NET chromatin meshwork highlights the crucial role of NETs as key mediators of thrombosis and its associated mortality (4–7). To avoid deleterious NET activity, the turnover of neutrophil-released DNA fibers is tightly controlled by the action of extracellular circulating DNases that degrade NETs (8). Endogenous blood-borne DNases play a pivotal role in regulating NET-mediated thrombotic vascular occlusions. Mice that are deficient in both DNase1 (D1) and DNase1-like 3 (D1L3) are susceptible to lethal NET-mediated thrombosis in experimental models of neutrophilia and sepsis (9). Similarly, a number of pathogenic bacteria secrete DNases that degrade NETs, thereby evading host defenses and facilitating bacterial dissemination (10).

A wide range of pathogen- and danger-associated molecular patterns (PAMPs, DAMPs) including lipopolysaccharide (LPS), immune complexes, reactive oxygen species (ROS) and cytokines (IL-1β), stimulate NETosis through nicotinamide adenine dinucleotide phosphate oxidase 2 (NOX2)-dependent or -independent pathways (11). NADPH oxidase-dependent NETosis, also referred to as “classic NETosis”, is typically initiated by microbial pathogens, lipopolysaccharide (LPS), N-formylmethionyl-leucyl-phenylalanine (fMLP), or phorbol-12-myristate-13-acetate (PMA). This process is dependent on protein kinase C (PKC) and cytosolic reactive oxygen species (ROS) (12). In contrast, NOX2-independent NET formation which is triggered by membrane permeabilizing calcium ionophore A23187 and ionomycin, is mediated by mitochondrial ROS (13, 14).

Over the past decade, significant progress has been made in understanding the molecular mechanisms underlying NETosis in specific contexts. In bacterial sepsis, the binding of damage-associated molecular patterns (DAMPs) and pathogen-associated molecular patterns (PAMPs) induces the dimerization of Toll-like receptors (TLRs) and the subsequent signaling via the attached adaptor protein, myeloid differentiation factor 88 (MYD88) (15, 16). Accordingly, *Myd88*-deficient mice are unresponsive to LPS (17) and protected from sepsis-stimulated intravascular NETosis (18, 19). Peptidyl arginine deiminase 4 (PAD4), which catalyzes the conversion of arginine residues to citrulline in a calcium-dependent manner (20), is another key enzyme involved in NET formation. Upon translocation to the nucleus, PAD4 citrullinates arginines in histones, thereby reducing their positive charge and electrostatic binding to DNA (21). Consequently, PAD4 promotes nucleosome disassembly and chromatin decondensation which ultimately results in the release of NETs. It was recently shown that the pore-forming protein gasdermin D (GSDMD) plays a crucial role in NET formation (22, 23). Upon canonical or non-canonical inflammasome activation, the N-terminal domain of GSDMD oligomerizes and forms pores at the plasma membrane. This results in cell swelling and subsequent membrane rupture, mediating NET release and pyroptosis, a form of lytic cell death (24, 25). However, other studies have demonstrated that PAD4 and GSDMD do not significantly contribute to NETosis under certain experimental conditions (26–28). Therefore, further investigation is required to elucidate the precise role of these proteins in NET formation during sepsis and sterile inflammation.

In this study, we compared the roles of MYD88, PAD4, and GSDMD in lethal vaso-occlusive NET formation using *in vivo* sepsis and neutrophilia models, triggered by LPS/*E. coli* infusion and granulocyte-colony stimulating factor (G-CSF or *Csf3*) transgene expression in *Dnase1/Dnase1l3*-deficient mice, respectively (9). *Myd88/D1/D1l3^−/−^* mice survived TLR-dependent LPS/*E.coli* challenge, however these mice succumbed to *Csf3*-induced neutrophilia. Unexpectedly, NET formation and lethal vascular occlusions readily proceeded in the absence of GSDMD and PAD4, both upon sepsis and neutrophilia challenge. Mortality from pulmonary embolism was similar in *Gsdmd/D1/D1l3^−/−^* and *Pad4/D1/D1l3^−/−^* mice as compared to *D1/D1l3^−/−^* mice. Our findings challenge the notion that GSDMD and PAD4 are indispensable for NET formation in sepsis- and neutrophilia-driven conditions. They also suggest that alternative pathways can sustain NETosis independently of the canonical mediators GSDMD and PAD4.

## RESULTS

### MYD88 is a critical mediator of sepsis- but not neutrophilia-induced NET formation

The two mouse models utilized in this study, sepsis and neutrophilia, are based on our previous work where we showed the necessity for secreted DNase1 and DNase1L3 to efficiently clear NETs from circulation (9). Mice lacking both *Dnase1* and *Dnase1l3* rapidly succumb to sepsis or neutrophilia challenge. Therefore, the objective was to assess the contribution of established regulators of NETosis in mitigating lethal NET-driven vascular occlusion in the context of sepsis and neutrophilia.

As a proof of concept for intravascular lethal NETosis, we initially confirmed that deficiency in MYD88 confers protection from TLR-mediated sepsis challenge. As anticipated, all (n = 7/7) *Myd88/D1/D1l3^−/−^* mice survived the sepsis challenge, as MYD88 signals downstream of TLRs. In contrast, >50% (n = 4/7) of *D1/D1l3^−/−^* mice died following infusion of LPS/*E. coli*. Wild-type (WT) mice with an intact NET-degrading capacity were similarly protected as *Myd88/D1/D1l3*^−/−^ mice (100% survival; n = 7/7; **Fig. 1A**). Progressive hypothermia with a median temperature decline of −2.51°C [interquartile range (IQR) −3.80 (25 % percentile) to 0.08 (75 % percentile)] preceded the death of *D1/D1l3^−/−^* animals. In contrast, all surviving mice recovered from hypothermia within five days post-challenge. In accordance with survival data, there was no significant difference in peripheral body temperature (median −0.18°C, IQR −0.75, 0.18) between sepsis-challenged *Myd88/D1/D1l3^−/−^* and WT mice on day 5 after treatment (**Fig. 1B**). *D1/D1l3^−/−^* mice exhibited a drop in body weight (median −3.09 g, IQR −3.78, −2.01), while *Myd88/D1/D1l3^−/−^* mice did not demonstrate a reduction in body weight throughout the 5 days (median −0.23 g, IQR −0.61, −0.01, *P* = 0.0096; **Fig. 1C**).

**Figure 1.**
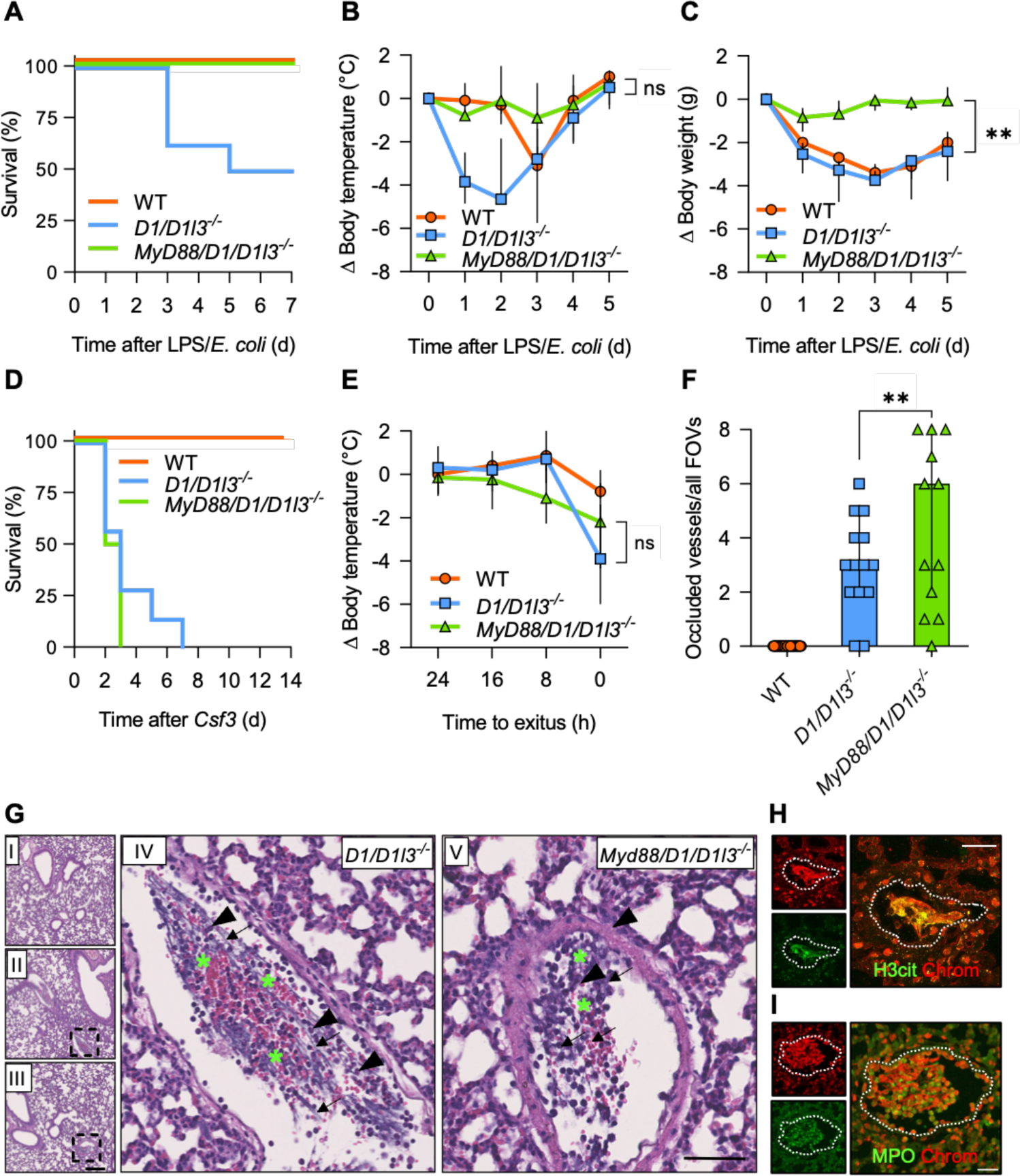
MYD88 is required for sepsis but not neutrophilia-driven intravascular NET formation. *Myd88/D1/D1l3^−/−^*, *D1/D1l3^−/−^*, and WT mice were i.p. injected with lipopolysaccharide (LPS, 3-times, 1 µg/g) and i.v. with 1.5 × 10^7^ heat-killed *E. coli* (**A-C**) or challenged with neutrophilia induced by hydrodynamic tail vein injection of a G-CSF-expressing plasmid (**D-I**). Survival (**A, D**), peripheral body temperature following LPS/*E. coli* infusions or *Csf3* expression (**B, E**) and body weight (**C**) of challenged mice is given. Histological analysis of *Myd88/D1/D1l3^−/−^ and D1/D1l3^−/−^* mice that ectopically expressed *Csf3* (**F-I**). Hematoxylin and Eosin (H & E) staining of lungs (**G**). A pulmonary blood vessel of *Csf3* over-expressing mice shows hematoxylin-rich clots, long DNA filaments (arrow) with entrapped erythrocytes (asterisk), and few leukocyte nuclei (arrowhead). Scale bars, 250 µm (overview) and 50 µm (zoomed-in view). Upper left overview **I** depicts an H & E-stained lung of a WT mouse. **II** is the overview of **IV**, a *D1/D1l3*^−/−^ mouse lung and **III** is the overview of **V**, a *Myd88/D1/D1l3^−/−^* mouse lung (**G**). Quantification of blood vessels in lungs occluded by hematoxylin-positive clots per field-of-view (FOV) in *Myd88/D1/D1l3^−/−^, D1/D1l3^−/−^* and WT mice, 5 FOVs/lung (**F**). Immunostaining of occluded blood vessels for chromatin (red) and the neutrophil markers H3cit and MPO (green, **H, I**). Scale bars, 25 µm. Representative images of four mice. Dotted lines indicate the vessel wall. Data is represented as median and interquartile range (IQR). (B, C, E) two-way ANOVA after 5 days or time point 0 h, (F) one-way ANOVA. * *P* < 0.05, ** *P* < 0.01, *** *P* < 0.001.

We next investigated whether deficiency in MYD88 could potentially rescue the lethal NET occlusive phenotype in a TLR-independent setting such as the neutrophilia mouse model. Neutrophilia can be induced by *Csf3* overexpression, which results in vascular NET formation in *D1/D1l3* deficient mice (9). As expected, *Myd88* deficiency did not provide protection from neutrophilia-induced vascular occlusions. All challenged *Myd88/D1/D1l3^−/−^* and *D1/D1l3^−/−^* animals died (n = 4/4 and n = 7/7, p>0.05; **Fig. 1D**). Mortality in both mouse lines was associated with a comparable magnitude of hypothermia (*Myd88/D1/D1l3^−/−^* median drop in body temperature −0.71°C, IQR −2.24, −0.35 *vs. D1/D1l3^−/−^* median 0.19°C, IQR −2.60, 0.60; **Fig. 1E**). Consistent with published data, none of the WT mice (n = 0/6) with normal D1/D1l3 expression succumbed to *Csf3* overexpression and only a minimal drop in body temperature (−0.80°C, median of n = 6) was triggered by neutrophilia. We analyzed pulmonary embolism by histological analysis. Hematoxylin and Eosin (H & E)-stained lung sections showed that the number of occluded vessels increased more than twofold in *Myd88/D1/D1l3^−/−^* mice compared to *D1/D1l3^−/−^* animals [median 6.00 occluded vessels/all field-of-views (FOVs), IQR 2.00, 8.00 *vs.* median 3.00 occluded vessels/FOV, IQR 2.00, 5.00, respectively, *P* = 0.0075; **Fig. 1F**]. Occluded vessel areas were rich in long DNA strands and the meshwork entrapped erythrocytes and leukocytes (**Fig. 1G**). Immunohistochemical analysis of NET biomarkers citrullinated histone H3 (H3cit), MPO, and chromatin provided confirmation that intravascular NET formation was the underlying mechanism responsible for pulmonary vascular occlusion in *Myd88/D1/D1l3^−/−^* and *D1/D1l3^−/−^* mice (**Figs. 1H, I**).

### Vascular occlusive NET formation occurs in the absence of GSDMD in sepsis and neutrophilia

To gain insight into the role of GSDMD-generated pores in the release of NETs into the extracellular space, we knocked out *Gsdmd* in *D1/D1l3*^−/−^ mice and challenged them in our two *in vivo* models. It is believed that the pore-forming protein GSDMD is required for NET formation. Both global *Gsdmd* gene ablation and pharmacological inhibition of GSDMD have been shown to interfere with NET formation in mouse models of lethal sepsis (29). We employed CRISPR/Cas9 technology to genetically target *Gsdmd* in *D1/D1l3^−/−^* mouse-derived zygotes and deleted the second coding exon (exon 3) of 193 base pairs (bp), resulting in a frameshift mutation (**Fig. 2A**). Quantitative reverse transcription-polymerase chain reaction (qRT-PCR) analysis confirmed mRNA expression of *Gsdmd* in tail biopsies of *Gsdmd/D1/D1l3*^−/−^ mice (**Fig. 2B**). Immunoblot analysis using antibodies that detect both the full-size protein at an apparent molecular weight of ∼ 53 kDa and its proteolytic N- and C-terminal fragments at ∼ 31 and ∼ 22 kDa, respectively, showed complete absence of GSDMD protein in *Gsdmd/D1/D1l3^−/−^* bone marrow cells (**Fig. 2C**). There was no obvious difference in development, fertility, body size and weight comparing *Gsdmd/D1/D1l3^−/−^* mice with age- and sex-matched *D1/D1l3^−/−^* or WT mice. Additionally, *Gsdmd/D1/D1l3^−/−^* animals exhibited a normal and healthy phenotype under baseline conditions.

**Figure 2.**
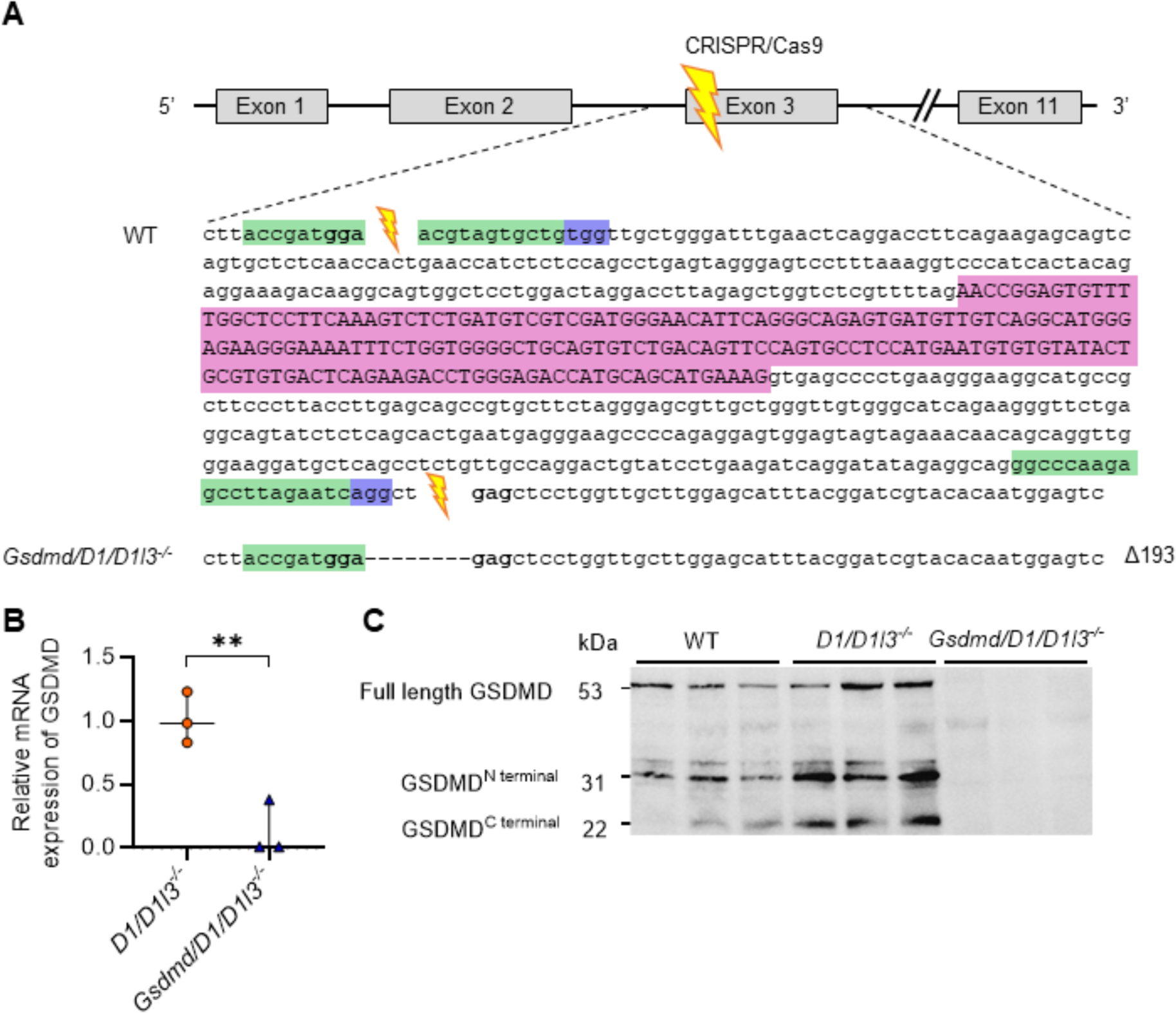
CRISPR/Cas9 gene disruption generates *Gsdmd^−/−^* mice. **(A)** Scheme showing the CRISPR/Cas9-mediated gene targeting strategy to generate *Gsdmd*^−/−^ mice. The sequence of the *Gsdmd* allele in WT and *Gsdmd^−/−^* mice with the intron-flanking sgRNA targeting exon 3 (2^nd^ coding exon) (green), target sequence (pink), and protospacer adjacent motif (PAM, purple) are depicted. Double-strand breakage by CRISPR/Cas9 followed by non-homologous end joining resulted in the deletion of 193 base pairs (bp) long exon 3. **(B)** *Gsdmd* mRNA expression from *Gsdmd/D1/D1l3^−/−^* and *D1/D1l3^−/−^* tail biopsies. (**C**) Lysed bone marrow cells isolated from *Gsdmd/D1/D1l3^−/−^*, *D1/D1l3^−/−^*, and WT mice were analyzed by western blotting using a polyclonal anti-GSDMD antibody. Equal protein loading of 20 µg/lane, n = 3 animals/genotype. Data is represented as median and interquartile range (IQR). (B) Student’s t-test. * *P* < 0.05, ** *P* < 0.01, *** *P* < 0.001.

We compared *Gsdmd/D1/D1l3^−/−^* and *D1/D1l3^−/−^* mice in our intravascular NET formation models induced by LPS/*E. coli* injection or *Csf3* overexpression. It was unexpected that over 80% of both *Gsdmd/D1/D1l3^−/−^* and *D1/D1l3^−/−^* mice succumbed within the first 48 hours following sepsis challenge (9/11 *vs.* 10/11). Furthermore, there was no significant difference in mortality in the absence of GSDMD (*P* > 0.05; **Fig. 3A**). In contrast, all WT control mice survived experimental sepsis (100%; n = 7/7). The number of thrombotic vascular occlusions was comparable between *Gsdmd/D1/D1l3^−/−^* and *D1/D1l3^−/−^* mice (1.00 occluded vessels/FOV, IQR 0.00, 2.00; **Fig. 3B**), as evidenced by H&E-stained lung sections. Multiple pulmonary vessels of both *Gsdmd/D1/D1l3^−/−^* and *D1/D1l3^−/−^* mice were occluded, consistent with lethal pulmonary embolisms in the two mouse lines (**Fig. 3C**). In contrast, there were virtually no thrombi observed in vessels of sepsis-challenged WT mice, confirming the critical role of circulating DNases in intravascular NET clearance. Immunohistochemical analyses of vascular thrombi showed an accumulation of neutrophils and filamentous DNA structures in both *Gsdmd/D1/D1l3^−/−^* and *D1/D1l3^−/−^* mice that also stained positive for H3cit, MPO and extracellular chromatin, suggestive of extensive NETosis (**Figs. 3D, E**).

**Figure 3.**
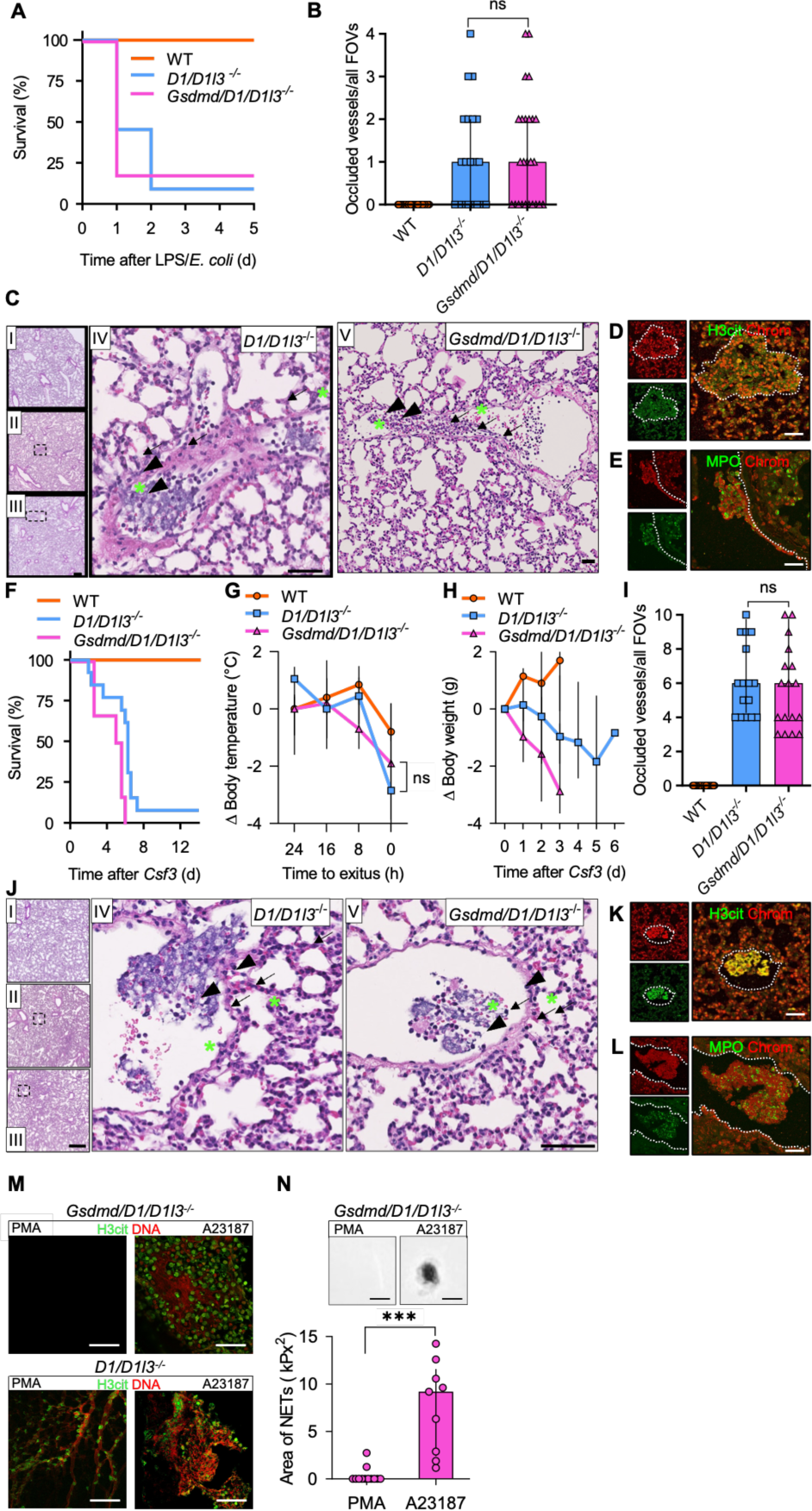
Vessel occlusive NET formation in sepsis and neutrophilia is independent of GSDMD. *Gsdmd/D1/D1l3^−/−^*, *D1/D1l3^−/−^*, and WT mice were challenged with LPS/*E. coli* (**A-E**) or *Csf3* overexpression (**F-L**) as shown in Fig. 1. Survival (**A, F**), change in peripheral body temperature 24 h before exitus (**G**), and change in body weight (**H**) of challenged mice. Histological analysis of *Gsdmd/D1/D1l3^−/−^* and *D1/D1l3^−/−^* mice following sepsis or *Csf3* expression challenge (**B-E, I-L**). H & E-stained lung sections revealed occluded pulmonary vessels. Hematoxylin-rich clots occluding vessels are made up of long DNA filaments (arrow) with entrapped erythrocytes (asterisk), and leukocytes (arrowhead). Scale bars, 250 µm (overview), **IV** 50 µm, and **V** 125 µm (zoomed-in view). Upper left overview **I** depicts a H & E-stained lung of a WT mouse. **II** is the overview of **IV**, a *D1/D1l3^−/−^* mouse lung and **III** is the overview of **V**, a *Gsdmd/D1/D1l3^−/−^* mouse lung. (**C, J**). Quantification of occluded blood vessels in lungs per field-of-view (FOV) in *Gsdmd/D1/D1l3^−/−^*, *D1/D1l3^−/−^* and WT mice, 5 FOVs/lung (**B, I**). Immunofluorescence images of occluded vessels stained for chromatin (red), H3cit, and MPO (green, **D, K** and **E, L**). Scale bars, 25 µm, representative images from four mice. Dotted lines indicate the vessel wall. Neutrophils were purified from neutrophilic *D1/D1l3^−/−^*, and *Gsdmd/D1/D1l3^−/−^* mouse blood, stimulated with PMA (10 nM) or A23187 (25 µM) and analyzed for NET generation at 4 h by immunohistochemistry staining of DNA (red) and H3cit (green). Representative images, n = 5 (**M**). Activated *Gsdmd/D1/D1l3^−/−^* mouse-derived neutrophils were visualized by brightfield microscopy and the surface area was quantified (**N**). Data is represented as median and interquartile range (IQR). (B, I) Kruskal-Wallis test, (G) two-way ANOVA at time point 0 h, (N) Mann-Whitney test. * *P* < 0.05, ** *P* < 0.01, *** *P* < 0.001.

In our second model of NET-mediated thrombosis driven by neutrophilia, we tested the unexpected redundancy of GSDMD for NETosis. As observed in septic conditions, the lethality of both *Gsdmd/D1/D1l3^−/−^* and *D1/D1l3^−/−^* mice was high, with >90% of animals dying within six days following neutrophilia challenge. Neutrophilia-driven mortality was not significantly different between the two mouse lines (6/6 *vs.* 12/13, *P* > 0.06; **Fig. 3F**). In contrast, all (n=6/6) WT mice survived the neutrophilia challenge. Severe and rapidly progressing hypothermia preceded the death of both *Gsdmd/D1/D1l3^−/−^* and *D1/D1l3^−/−^* mice (median −0.43°C, IQR −3.10, −0.32 *vs.* 0.35°C, IQR −1.86, 0.78, ns), whereas no significant change in peripheral body temperature was noted in challenged WT mice (median 0.16°C, IQR −0.36, 0.60; **Fig. 3G**). Reduction in body weight was not significantly different in *Gsdmd/D1/D1l3^−/−^* and *D1/D1l3^−/−^* mice either (−1.16 g, IQR −2.13, −0.21 *vs.* −0.57 g, IQR −0.88, −0.03, p>0.05; **Fig. 3H**). In line with the sepsis model, clot burden in pulmonary vessels was similar in challenged *Gsdmd/D1/D1l3^−/−^* and *D1/D1l3^−/−^* mice (6.00 occluded vessels/FOV, IQR 4.00, 9.75 *vs.* 6.00 occluded vessels/FOV, IQR 4.25, 9.75; **Fig. 3I**). Occlusions were rich in NETs as revealed by H&E staining of lung sections (**Fig. 3J**) and immunohistochemistry signals for chromatin, H3cit and MPO (**Figs. 3K, L**).

To better understand GSDMD functions in NETosis, we stimulated isolated neutrophils from *Gsdmd/D1/D1l3^−/−^* mice with two distinct activators, PMA and the Ca^2+^ ionophore A23187, which differ in downstream pathways leading to NET formation (30). Ten nM PMA was inactive in triggering NETosis in *Gsdmd/D1/D1l3^−/−^* neutrophils, but readily activated NET formation in *D1/D1l3^−/−^* neutrophils seen by immunofluorescent staining for MPO, H3cit and DAPI (**Fig. 3M, left panels**). Quantitation of PMA-induced NETs confirmed the virtual absence of NETosis in the absence of GSDMD (DNA-covered area in brightfield images of 0.0 kpx², IQR 0.0, 0.0 *vs.* 2.20 kpx^2^, IQR 1.51, 2.89 in *D1/D1l3^−/−^* neutrophils [not shown]; **Fig. 3N, left column**). *Gsdmd/D1/D1l3^−/−^* neutrophils could form NETs upon A23187 treatment as revealed by immunofluorescence staining for H3cit/DNA NET biomarkers (**Fig. 3M, right panels**), with a median NET clot area of 9.18 kpx² (IQR 2.39, 11.58; **Fig. 3N, right column**). In conclusion, these data show that GSDMD is essential for PMA-induced NET formation *in vitro* but not for NETosis associated with sepsis or neutrophilia *in vivo*.

### PAD4 is not a requisite factor for the formation of occlusive NETs in sepsis and neutrophilia

To investigate the necessity of histone citrullination in the process of chromatin decondensation during NET formation, we crossed *Pad4^−/−^* mice with *D1/D1l3^−/−^* mice. *Pad4/D1/D1l3^−/−^* mice and *D1*/*D1l3^−/−^* controls were similarly susceptible to LPS/*E.coli* infusions with 9/9 and 7/9 of the challenged mice dying within four days, respectively (**Fig. 4A**). In contrast, all WT mice (n=7/7) survived the sepsis. In line with survival, *Pad4/D1/D1l3^−/−^* mice exhibited a rapid drop in body weight, whereas surviving *D1*/*D1l3^−/−^* and WT control mice recovered after four days (*Pad4/D1/D1l3^−/−^*-median −3.85 g, IQR −7.08, −0.80, *D1/D1l3^−/−^* - median −3.16°C, IQR −4.02, −1.83, WT: median −2.28 g, IQR −3.00, −1.38; **Fig. 4B**). The numbers of occluded pulmonary vessels were not significantly different between *Pad4/D1/D1l3^−/−^* and *D1/D1l3^−/−^* animals (median 2.00 occluded vessels/all FOVs, IQR 0.00, 3.00 *vs.* median 3.00 occluded vessels/all FOVs, IQR 1.00, 4.00, respectively). In contrast, vascular occlusions were virtually absent in WT mice (**Fig. 4C**). Occlusive thrombi in the two lines appeared morphologically similar and were rich in DNA filaments and polynucleated cells (**Fig. 4D**). *Pad4/D1/D1l3^−/−^* lung tissue sections stained positive for MPO and chromatin in immunohistochemistry. As expected, pulmonary thrombi in *Pad4/D1/D1l3^−/−^* animals were defective in histone citrullination, seen by loss of H3cit signal in immunohistochemistry (**Figs. 4E, F**).

**Figure 4.**
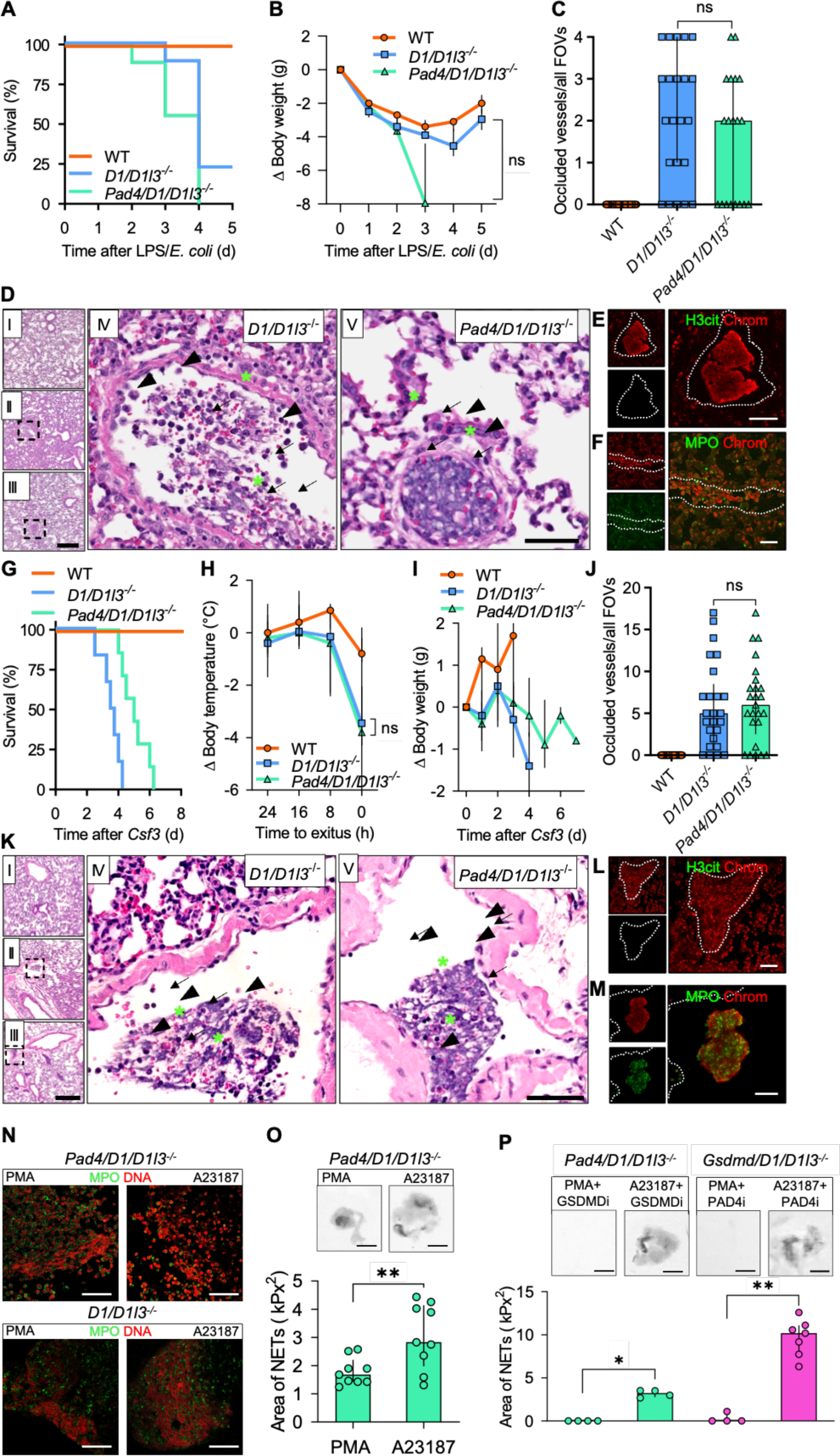
PAD4 deficiency allows for intravascular NET formation during sepsis and neutrophilia. *Pad4/D1/D1l3^−/−^*, *D1/D1l3^−/−^*, and WT mice were challenged by LPS/*E. coli* (**A-F**) or *Csf3* overexpression (**G-M**) as given in Figs. 1. and 3. Survival (**A, G**), change in body weight (**B, I**), and temperature 24 h before exitus (**H**). H & E-stained lung sections showed occluded vessels with clots comprised of DNA filaments (arrow), erythrocytes (asterisk), and leukocytes (arrowhead). Scale bars, 250 µm (overview) and 50 µm (zoomed-in view). Upper left overview **I** depicts a H & E-stained lung of a WT mouse. **II** is the overview of **IV**, a *D1/D1l3*^−/−^ mouse lung and **III** is the overview of **V**, a *Pad4/D1/D1l3^−/−^* mouse lung (**D, K**). Number of occluded blood vessels in lungs per field-of-view (FOV) in *Pad4/D1/D1l3^−/−^ and D1/D1l3^−/−^* mice, 5 FOVs/lung (**C, J**). Immunofluorescence images of vessel-occluding aggregates show positive staining for chromatin (red) with H3cit or MPO (green). Scale bars, 25 µm, n = 4. Dotted lines indicate the vessel wall (**E, L** and **F, M**). Neutrophils were purified from neutrophilic *D1/D1l3^−/−^*, and *Pad4/D1/D1l3^−/−^* mouse blood, stimulated with PMA (10 nM) or A23187 (25 µM) and analyzed for NET generation at 4 h by immunohistochemistry staining of DNA (red) and MPO (green). Representative images, n = 5 (**N**). Activated *Pad4/D1/D1l3^−/−^* mouse-derived neutrophils were visualized by brightfield microscopy and the NET area was quantified from these microscopic images (**O**). Neutrophils from *Pad4/D1/D1l3^−/−^* and *Gsdmd/D1/D1l3^−/−^* mouse blood were treated with the GSDMD inhibitor necrosulfonamide (GSDMDi, 10 µM) and the PAD4 inhibitor GSK484 (PAD4i, 5 µM) in parallel to the addition of the activators PMA or A23187, as shown by brightfield images that were quantified respectively (**P**). Data is represented as median and interquartile range (IQR). (B, C, J) Kruskal-Wallis test after 3 days (B), (H) two-way ANOVA at time point 0 h, (O) Student’s t-test, (P) Mann-Whitney test.* *P* < 0.05, ** *P* < 0.01, *** *P* < 0.001.

Spontaneous NET formation in the neutrophilia model also demonstrated that all *Pad4/D1/D1l3^−/−^* (n=7/7) and *D1/D1l3^−/−^*(n=6/6) mice died within six days after *Csf3* overexpression, while WT mice (n=6/6) survived the challenge (**Fig. 4G**). Concomitantly, a significant drop in body temperature preceded mortality by 8 h and was similarly reduced in *Pad4/D1/D1l3^−/−^* and *D1/D1l3^−/−^* mice (median −1.01°C, IQR −2.31, 0.12 *vs.* median −0.33°C, IQR −3.07, 0.27), while temperature remained stable in WT mice (median 0.16°C, IQR −0.36, 0.60; **Fig. 4H**). *Pad4/D1/D1l3^−/−^* and *D1/D1l3^−/−^* mice exhibited a reduction in body weight (median - 0.11 g, IQR −0.60, 0.14, *vs.* median −0.10, IQR −0.85, 0.03), in contrast to WT animals [(WT: median 0.58 g, IQR 0.10, 1.00) **Fig. 4I**]. Vascular occlusions in H & E-stained lung sections showed that the number of thrombi in *Pad4/D1/D1l3^−/−^* was not significantly different compared to *D1/D1l3^−/−^* control mice (median 6.00 occluded vessels/ FOVs, IQR 2.50, 8.50 *vs.* median 5.00 occluded vessels/ FOVs, IQR 1.50, 8.50, respectively). In contrast, WT mice did not develop any thrombi and sections were effectively free of occlusions (**Figs. 4J, K**). Immunohistochemical analysis of lung sections indicated that pulmonary occlusions contained NETs and stained for chromatin and MPO but not H3cit in *Pad4/D1/D1l3^−/−^* mice (**Figs. 4L, M**). These findings collectively indicate that *Pad4/D1/D1l3^−/−^* mice succumbed to vascular NET-derived occlusions in sepsis and neutrophilia.

Next, we compared NET formation in neutrophils isolated from *Pad4/D1/D1l3^−/−^* mice by stimulating cells *ex vivo* with PMA and A23187. Neutrophils from *Pad4/D1/D1l3^−/−^* and *D1/D1l3^−/−^* mice formed NETs after 4 h of stimulation with PMA and A23187, as shown by immunohistochemistry staining for DNA and MPO (**Fig. 4N**). Quantitation of NET clots revealed an increase in the clot area when *Pad4/D1/D1l3^−/−^* neutrophils were stimulated with A23187 in comparison to PMA (PMA: median clot area of 1.68 kpx², IQR 1.43, 2.21; *vs.* A23187: 2.83 kpx², IQR 1.97, 4.14, *P* = 0.0086; **Fig. 4O**).

To test for potential mutual compensation of PAD4 and GSDMD functions in NET formation, we pharmacologically inhibited GSDMD (GSDMDi, necrosulfonamide, NSA, 10 µM) or PAD4 (PAD4i, GSK484, 5 µM) in *Pad4*-or *Gsdmd*-deficient mouse neutrophils, respectively. Targeting GSDMD abolished NET formation in PMA-but not A23187-activated PAD4 deficient neutrophils (PMA+GSDMDi: median 0.0 kpx², IQR 0.0, 0.0; *vs.* A23187+GSDMDi: median 3.24 kpx², IQR 2.76, 3.52, *P* = 0.0286). Pharmacologic interference with PAD4 activity using GSK484 (5 µM), did not significantly reduce A23187-activated NET formation in *Gsdmd*-deficient neutrophils (A23187+PAD4i: median 10.18 kpx², IQR 7.73, 11.10; **Fig. 4P**).

## DISCUSSION

Dysregulated or excessive NETosis contributes to thromboinflammatory disease states and PAD4 and GSDMD were previously shown to be critical for NETosis *in vivo* (29, 31, 32). Here, we evaluated the role of PAD4 and GSDMD in two distinct models of lethal vessel occlusive NET formation (22, 23). Our findings indicate that both GSDMD (**Fig. 3**) and PAD4 (**Fig. 4**) are not essential for intravascular NET formation in response to *S. typhimurium* LPS/*E. coli*-triggered sepsis and *Csf3* overexpression-driven neutrophilia.

The TLR-MYD88 pathway activates the mitogen-activated protein kinase (MAPK) and nuclear factor kappa-light-chain-enhancer of activated B cells (NF-κB) signaling pathways which lead to the production of pro-inflammatory interleukins and cytokines. These in turn activate neutrophils and induce NET formation (33–35). We found that *Myd88/D1/D1l3^−/−^* mice were protected from sepsis-but not neutrophilia-stimulated NETosis (**Fig. 1**) indicating that our NET-driven thrombosis model is suitable for assessing non-sterile NET formation. Indeed, bacterial PAMPs and DAMPs stimulate TLR signaling through MYD88, thereby rendering *Myd88/D1/D1l3^−/−^* mice resistant to sepsis but susceptible to neutrophilia (**Fig. 1**).

The results of recent studies have been inconclusive regarding the importance of GSDMD for NET formation (22, 23, 28, 29, 36, 37). GSDMD deficiency contributes to survival following *E. coli* O111:B4 LPS-stimulated endotoxemia (38–40) and in cecal ligation and puncture (CLP)-induced polybacterial sepsis models (41). Consistently, downstream markers of GSDMD activation such as IL-1α and IL-1β, correlated with disseminated intravascular coagulation (DIC)-scores in sepsis patients, supporting a role for GSDMD in bacterial infection and likely NETosis (42). In line with these findings, genetic or pharmacologic inhibition of GSDMD abrogated NET formation and improved survival in CLP sepsis and endotoxemia models (29). In the present study, we did not observe that GSDMD deficiency significantly impacted NETosis and survival in our models. The discrepancy between our findings and those of other studies may be attributed to the use of different models and stimuli to induce NETosis. CLP-triggered sepsis and LPS-induced endotoxemia (single bolus of 10 µg/g) may induce a more acute reaction *in vivo* compared to the sepsis model that we employed which entails repeated i.p. injections of low-dose LPS/*E.coli* over several days. Indeed, when we established our sepsis model, daily i.p. injections of low dose LPS (1 μg/g) in *D1/D1l3^−/−^* mice resulted in vascular occlusions of only a small portion of pulmonary vessels (approximately 20%) and all mice survived the procedure (9). The addition of heat-killed *E. coli* with the third LPS injection resulted in an increased mortality rate (9). In accordance with our findings, Liu and colleagues reported that mice with a neutrophil-specific conditional deletion of GSDMD (*Gsdmd*^flox/flox^MRP-Cre^+^) were also not protected from CLP-induced sepsis. Similar to our global *Gsdmd/D1/D1l3^−/−^* mice, neutrophil-specific *Gsdmd* deficiency did not provide protection from sepsis and tissue damage, whereas plasma levels of proinflammatory cytokines and NET formation were similar to WT mice (28). Together, these data indicate that the kinetics and magnitude of a stimulus in sepsis-induced NET formation likely triggers diverse molecular mechanisms and show that GSDMD-independent pathways, that are yet to be identified, are required for NETosis in sepsis.

For example, mixed lineage kinase domain-like (MLKL), another pore-forming protein with similarities to GSDMD, has been demonstrated to induce NET extrusion into the extracellular space in mice and humans (43–45). While GSDMD cleavage induces NETosis and pyroptosis (23), MLKL has been linked to necroptosis, driven by ligand binding to TNF family death domain receptors, pattern recognition receptors, and virus sensors (46, 47). The partial compensation for GSDMD and MLKL functions is further evidenced by the significantly enhanced protection from polymicrobial sepsis observed in mice with combined deficiency in Mlkl and Gsdmd, as compared to the protection observed in either *Mlkl^−/−^* and *Gsdmd^−/−^* mice (48). Moreover, elevated plasma levels of MLKL correlate with the severity and outcomes in critically ill sepsis patients (48, 49). Therefore, under specific circumstances, the actions of other pore-forming enzymes, such as MLKL, may become crucial and serve to compensate for GSDMD in the promotion of NETosis. Potential alternative mediators that may substitute for GSDMD could include histone modifications such as a reduction in positive charge that would limit binding to DNA, or defective chromatin packing which would result in increased entropic pressure, chromatin swelling, and plasma membrane rupture, independently of GSDMD activity (50).

The role of PAD4 in NET formation remains a topic of contention (51–54). Previous studies have demonstrated that Pad4 deficiency is not a prerequisite for NET formation and does not influence the outcomes of murine bacterial sepsis induced by *K. pneumoniae* (26) or *C. albicans* (27). Indeed, *Pad4^−/−^* and WT mice infected with *K. pneumoniae* showed similar NET structures, cell-free DNA levels, bacterial growth, lung inflammation, and organ damage (26). Likewise, *Pad4^−/−^* mice infused with *C. albicans* exhibited comparable NET formation and antifungal activity compared to WT mice, despite a loss of histone citrullination (27). PAD4-independent NET formation has been identified in amoeba infection (55), antibody-mediated arthritis (56) and influenza A infection (51), while PAD4 is required for NET formation in lupus (57, 58), gallstone formation (59), bacterial infection with *S. flexneri, P. aeruginosa* and *S. aureus* (31, 60), and arterial and venous thrombosis (32, 61). Furthermore, murine neutrophils stimulated with calcium ionophores underwent NETosis irrespective of the presence or absence of the PAD4 inhibitor, Cl-amidine (27). We found that *Pad4*-deficient neutrophils retained their capacity to release NETs *ex vivo* following stimulation with PMA or A23187 (**Fig. 4**). An increase in intracellular calcium concentration [Ca^2+^]_i_ initiates the formation of non-selective mitochondrial pores (mPTP) and mitochondrial reactive oxygen species (mtROS) which provide an alternative pathway for NET formation independently of PAD4 activation (14, 62). Indeed, ROS and mtROS stimulate the release of neutrophil granule proteins, including MPO and NE, which translocate to the nucleus, where they modify chromatin in a PAD4-independent manner (63). These alternative pathways for chromatin decondensation driven by NE-mediated histone and laminin breakdown, may compensate for PAD4 and allow for NET formation in *Pad4 ^−/−^* mice (64). Collectively, our data demonstrate that GSDMD and PAD4 are not essential for in vivo NET formation in our models of sepsis or neutrophilia. This suggests that other, as yet unidentified mechanisms may serve to compensate for deficiencies in PAD4 and GSDMD during NET formation.

## MATERIALS AND METHODS

### Experimental animals

All mice were bred on a C57BL/6 genetic background for >10 generations. Drs. Markus Napirei (67) and Ryushin Mizuta (68) provided *Dnase1*^−/−^ and *Dnase1l3^−/−^* mice, respectively. We crossed *Dnase1*^−/−^ and *Dnase1l3^−/−^* mice to generate *Dnase1/Dnase1l3^−/−^* mice (9). *Myd88^−/−^* mice were purchased from Jackson Laboratory (B6.129P2(SJL)-Myd88tm1.1Defr/J; strain 009088) and crossed with *Dnase1/Dnase1l3^−/−^* mice. *Pad4^−/−^* mice were provided by Dr. Kerri A. Mowen (51) and crossed to a *Dnase1/Dnase1l3^−/−^* background. The *Dnase1/Dnase1l3/Gsdmd^−/−^* mice were generated by Dr. Irm Hermans-Borgmeyer, Transgenic Mouse Unit, University Medical Center Hamburg-Eppendorf using CRISPR/Cas9 technology, which is described below. DNA genotyping confirmed successful gene targeting.

### Murine blood, plasma, and tissue collection

Blood was collected from the retro-orbital sinus into 200 µl EDTA tubes (KABE Labortechnik, GK 150). Plasma was collected after centrifugation at 3,000 x g for 15 min and stored in aliquots at −20°C until further use. For organ analysis, mice were intracardially perfused with 20 ml PBS. Organs were collected and fixed for 24 h in 4% formaldehyde at 4°C. Fixed organs were embedded in paraffin for tissue sectioning.

### Neutrophil isolation from mouse blood

One ml of EDTA full blood was diluted in PBS containing 1% (w/v) bovine serum albumin (BSA) and 15 mM EDTA and fractionated on a discontinuous sucrose gradient. 3 ml of sterile-filtered sucrose 1.119 g/ml was added to the bottom of a 15 ml conical polypropylene centrifuge tube. Three ml of sterile-filtered sucrose 1.077 g/ml were added on top of the 1.119 g/ml layer. Thereafter, 6 ml of diluted blood was layered on top of the 1.077 g/ml layer and centrifuged at 700 x g for 30 min at room temperature without brake. Neutrophil isolation was continued using the negative selection method according to the manufacturer’s protocol of the neutrophil isolation kit (MACS Miltenyi, 130-097-658).

### *Ex vivo* generation of NETs

Mouse neutrophils were seeded at 10^7^ cells/ml in DMEM supplemented with 2% BSA. Neutrophils were stimulated with 25 µM calcium ionophore A23187 (Sigma, C5149) or 10 nM phorbol-12-myristate-13-acetate (PMA, Sigma, P8139) for 4 h, in the presence or absence of the necroptosis/pyroptosis inhibitor necrosulfonamide (NSA, 10 µM, Selleck, S8251) or the PAD4 inhibitor GSK484 (5 µM, Sigma, SML1658) for 30 min at 37°C, 5% CO_2_ with humidity and gentle shaking. NET degradation was controlled by treating NETs with 10 U/ml recombinant DNase1 (dornase alpha, Roche) for 1 h at 37°C, which resulted in a complete degradation of the formed NETs. NET clots were fixed with 2% formaldehyde overnight at 4°C. Fixed NET clots were embedded in paraffin and sections were analyzed for NET markers as described below. The area of NETs (in kPx) was quantified using the polygon setting in ImageJ.

### Immunofluorescence staining

Paraffin-embedded sections of *ex vivo* produced NETs or lung tissue were de-paraffinized, rehydrated, and subjected to heat-induced antigen retrieval for 25 min at 100°C in citrate buffer (10 mM sodium citrate, 0.1% Tween, pH 6). Slides were blocked for 1 h with PBS supplemented with 3% BSA and 0.3% Triton X-100. Primary antibodies were incubated overnight at 4°C, washed 3 times with PBST, and subsequently incubated with secondary antibodies for 1 h. Next, sections were washed, quenched using Sudan Black (0.1% in 70% ethanol) and mounted. The following primary antibodies were used: anti-MPO (2 µg/ml, Dako, A0398), anti-citrullinated histone 3 (2 µg/ml, Abcam, ab5103), and anti-histone H2A/H2B/DNA-complex (anti-chromatin, 5 µg/ml, Davids Biotechnologie GmbH, (4)). Secondary antibodies conjugated to AlexaFluor-488 and −594 were obtained from Jackson ImmunoResearch (all donkey, used in 1:200 dilution). Images were acquired with an inverted fluorescence microscope (Axiovert 200M, Zeiss) or an inverted confocal microscope (Leica TCS SP8, Leica).

### Development of *Gsdmd*-deficient mice

Three sgRNAs (5’ from target: ACC GAT GGA ACG TAG TGC TG; central in target: AAA GTC TCT GAT GTC GTC GA; 3’ from target: GGC CCA AGA GCC TTA GAA TC) flanking and targeting exon 3 (2^nd^ coding exon) of *gasdermin d* (Q9D8T2) were designed and templates for transcription were derived by PCR using Q5-Polymerase (Biolabs, M0491S). Transcription was performed using the HiScribeT7 kit (Biolabs, E20140S) with subsequent purification of the transcripts with the MEGAClear^TM^ kit (Fisher Scientific, AM1908), both according to the manufacturer’s instructions. Electroporation into single cell stage embryos derived from *Dnase1/Dnase1l3^−/−^* mice using 300 ng/µl of each sgRNA and 500 ng/ml Cas9 protein (Integrated DNA Technologies, 1081058) in OptiMEM medium was performed with a NEPA 21 electroporator. Settings used as previously described by Remy *et al.* (69). Electroporated embryos were implanted into F1 foster mothers (C57BL6/J x CBA). Resulting offspring was genotyped using PCR and mice lacking exon 3 were crossed to *Dnase1/Dnase1l3^−/−^ mice*, genotyped by PCR and sequencing of the 5’ and 3’ neighboring region of the deletion was performed.

### Genotyping PCRs

DNA from tail biopsies was isolated using the blackPREP Rodent Tail DNA Kit (IST Innuscreen, 845-BP-0010250) according to the manufacturer’s protocol. DNA templates were amplified on the PTC-200 thermal cycler (MJ Research) using the KAPA2G Fast HotStart Genotyping Mix (Roche, KK5621) and the respective PCR primers (Eurofins, Germany) for all the mouse lines except for *Gsdmd/D1/D1l3^−/−^* mice. DNA from *Gsdmd/D1/D1l3^−/−^* mouse tails was amplified using a genotyping mix containing a dNTP mix (10 mM, Qiagen, 201901), *Taq* DNA polymerase (250 U, Qiagen, 201203), and the respective primers. Amplified DNA fragments were visualized by agarose gel electrophoresis. The following PCR primers were used to amplify the genes of interest.

*Dnase1* KO CRLSV40pA_F primer: 5’ CAC TGC ATT CTA GTT GTG GTT TGT C 3’

*Dnase1* KO tm1_mR1 primer: 5’ GAG GCA GGA CTT AAT ACA CAA ACA G 3’

*Dnase1-like 3* KO G3 primer: 5’ GGG CCA GCT CAT TCC TCC ACT C 3’

*Dnase1-like 3* KO R7 primer: 5’ CAC TCC TGG GCT TCT TGA TGG TCA G 3’

*Myd88* oIMR9481 primer: 5’ GTT GTG TGT GTC CGA CCG T 3’

*Myd88* oIMR9482 primer: 5’ GTC AGA AAC AAC CAC CAC CAT GC 3’

*Myd88* 9335 primer: 5’ CCA CCC TTG ATG ACC CCC TA 3’

*Pad4* 3loxF primer: 5’ AGT CCA GCT GAC CCT GAA C 3’

*Pad4* 3loxR primer: 5’ CTA AGA GTG TTC TTG CCA CAA G 3’

*Pad4* 5loxF primer: 5’ CAG GAG GTG TAC GTG TGC A 3’

*Gsdmd* F primer: 5‘ GTT CCC TCC AGC CCT ACT TGC TC 3’

*Gsdmd* Rev primer: 5‘ GAG AAG TGG ACA CTC GTG CCT GTG 3’

### RT-PCR

RNA from tail biopsies was isolated using the RNeasy Mini Kit (74104, Qiagen) according to the manufacturer’s protocol. Reverse transcription of RNA into cDNA was performed using the SuperScript IV VILO Master Mix (Thermo Fisher Scientific, 11756050). The cDNA was then quantified with the *Gsdmd* gene probe for mice (Taqman Gsdmd Mm00509958_m1, Thermo Fisher Scientific, 4331182) and run on the StepOnePlus Real-Time PCR System (Applied Biosystems).

### Isolation of bone marrow cells

From each mouse, the bone marrow cells were flushed from both femurs and tibias with ice-cold PBS. The cells were resuspended with ice-cold PBS and centrifuged at 2,000 x g for 5 min at 4°C, this step was repeated twice. The cell pellets were then lysed with RIPA lysis buffer (Thermo Fisher Scientific, 89901) supplemented with Complete Protease Inhibitor Cocktail (Roche, 4693132001) for 30 min at 4°C with agitation. Cell lysates were centrifuged at 16,000 x g for 20 min, supernatants were collected, and protein concentrations were quantified by the BCA protein assay kit (Thermo Fisher Scientific, 23225).

### Immunoblotting

Equal amounts of bone marrow cell lysates underwent electrophoresis on a gradient SDS-PAGE gel (Criterion XT precast gel, 4-12%, BioRad, 3450123). Proteins were transferred onto nitrocellulose membranes and after blocking with Intercept (TBS) Blocking Buffer (Li-Cor, 927-60001) incubated with GSDMD antibody (1:1,000, Abcam, ab219800) overnight at 4°C, followed by a 1 h incubation with secondary goat anti-rabbit IgG (1:10,000, Abcam, ab6721). Antibodies were diluted in blocking buffer supplemented with 0.2% Tween-20. The results were visualized using the ChemiDoc Imager (Bio-Rad).

### Production of *Csf3* transgenic mice

Hydrodynamic tail vein injection of the pLIVE plasmid (Mirus Bio, USA) was used to express proteins in a long-lasting and hepatocyte-specific manner in mice, as described before (9). In brief, 50 µg of plasmid were diluted in 0.9% saline in a volume equivalent to 10% of the body weight of the mouse. Mice were anaesthetized with isoflurane before the plasmid solution was administered intravenously within 5 – 8 seconds via the tail vein. In the rare case that a mouse did not fully recover from the injection within the first 24 h, it was excluded from the study.

### Neutrophilia

Mice at 8-12 weeks of age were injected with 50 μg of the pLIVE plasmid containing *Csf3* to induce G-CSF expression and neutrophilia using hydrodynamic tail vein injections (9). Neutrophils counts (CD11b^+^ Ly6G^+^ cells) increased to up to 20 × 10^3^ cells/µl at 14 days after *Csf3* treatment (9). Mice were euthanized and considered “non-survivors” when showing defined signs of distress (no spontaneous movement, closed eyes, occasional gasping) accompanied by hematuria and rapid and severe hypothermia, defined as a decrease in body temperature of ≥ 4°C compared to the body temperature before the plasmid injection. The body temperature of the mice was measured in the perianal area using a contactless infrared thermometer, and body weight was determined with a scale. Blood and organs were collected at the time of euthanasia. Mouse experiments were performed according to national guidelines for animal care and were covered by animal proposals 69/16 and 143/15, approved by the Ministry for Health and Consumer Protection in Hamburg, Germany.

### Sepsis

Mice at 6-12 weeks of age received daily intraperitoneal injections of 1 µg/g LPS from *Salmonella enterica* serotype *thyphimurium* (Sigma, L6511, lot 025M4042V, strain ATCC 7823) in 0.9% saline, along with an intravenous injection of 1.5 × 10^7^ heat-killed *Escherichia coli* (*E. coli*)/g body weight (Perkin Elmer, XEN 14 derived from the parental strain *E. coli* WS2572) on the third day of LPS treatment (9). *E. coli* was previously prepared by growing the bacterial cells overnight in lysogeny broth medium supplemented with 50 µg/ml kanamycin, then centrifuged at 4,000 x g for 10 min, washed and resuspended in PBS. Next, the bacteria were heat-inactivated by incubation at 70°C for 15 min and stored at −20°C until further use. Survival time indicates the time after the intravenous injection of *E. coli*. Mice were euthanized and considered “non-survivors” when showing defined signs of distress combined with severe hypothermia and hematuria. Blood and organs were collected at the time of euthanasia. Mouse experiments were performed according to national guidelines for animal care and were covered by the animal proposal 22/17, approved by the Ministry for Health and Consumer Protection in Hamburg, Germany.

### Statistical evaluation

Statistical analyses of the data included Student’s t-test, Mann-Whitney test, one-way ANOVA followed by Tukey’s test, or Kruskal-Wallis followed by Dunn’s test, depending on the number of analyzed groups and data distribution. Results were considered significant at *P* < 0.05. All statistical analyses were performed with Prism version 8.2.0 (GraphPad, USA). In the case of multiple survival curves at 100%, they were optically separated for clarification.

## CONFLICT OF INTEREST

T.A.F. is co-founder and CEO of Neutrolis Inc. All other authors have nothing to declare.

## AUTHOR CONTRIBUTIONS

H.E., C.R., M.D., and J.G. performed experiments. I.H.B. and U.B. generated CRISPR/Cas9 mice. M.B., R.K.M, M.G., M.F., S.W.S. and R.P. conceived and discussed data and figures.

H.E. and C.R. generated figures. T.A.F. and T.R. conceived, supervised, and directed the study and provided funding. H.E., C.R., E.X.S. and T.R. wrote and edited the manuscript.

## FUNDING

This study was supported by the P06/KFO306 and INST 152/876-1 FUGG grants of the DFG (to T.R.).

## ACKNOWLEDGEMENTS

We acknowledge the UKE Microscopy Imaging Facility UMIF (DFG Research Infrastructure Portal: RI_00489) and the UKE Mouse Pathology Core Facility.

